# Dissociable roles of the mPFC-to-VTA Pathway in the control of iImpulsive Action and Risk-Related Decision-Making in Roman High- And Low-Avoidance Rats

**DOI:** 10.1101/2024.03.13.584562

**Authors:** Ginna Urueña-Méndez, Chloé Arrondeau, Florian Marchessaux, Raphaël Goutaudier, Nathalie Ginovart

**Affiliations:** Faculty of Medicine, Department of Psychiatry, University of Geneva, Geneva, Switzerland; Faculty of Medicine, Department of Basic Neuroscience, University of Geneva, Geneva, Switzerland

**Keywords:** Impulsive action, risk-related decision-making, mPFC-to-VTA pathway, [^18^F]-FDG

## Abstract

Impulsivity is a multidimensional trait associated with various psychiatric disorders including drug abuse. Impulsivity facets, such as impulsive action and risk-related decision-making (RDM), have been associated with reduced frontocortical activity and alterations in dopamine function in the ventral tegmental area (VTA). However, despite direct projections from the medial prefrontal cortex (mPFC) to the VTA, the specific role of the mPFC-to-VTA pathway in the control of impulsive behaviors remains unexplored. Here, we used Positron Emission Tomography with [^18^F]-Fluorodeoxyglucose to evaluate brain metabolic activity in Roman High-(RHA) and Low-avoidance (RLA) rats, which exhibit innate differences in impulsivity. Notably, we used a viral-based intersectional chemogenetic strategy to isolate, for the first time, the role of the mPFC-to-VTA pathway in controlling impulsive behaviors. We selectively activated the mPFC-to-VTA pathway in RHAs and inhibited it in RLAs, and assessed the effects on impulsive action and RDM in the rat gambling task. Our results showed that RHA rats displayed higher impulsive action, less optimal decision-making, and lower cortical activity than RLA rats at baseline. Chemogenetic activation of the mPFC-to-VTA pathway reduced impulsive action in RHAs, whereas chemogenetic inhibition had the opposite effect in RLAs. However, these manipulations did not affect RDM. Thus, by specifically and bidirectionally targeting the mPFC-to-VTA pathway in a phenotype-dependent way, we were able to revert innate patterns of impulsive action, but not RDM. Our findings suggest a dissociable role of the mPFC-to-VTA pathway in impulsive action and RDM, highlighting its potential as a target for investigating impulsivity-related disorders.

## Introduction

Impulsivity is a multifaceted personality trait characterized by a tendency to respond prematurely, i.e., impulsive action, and to make decisions without thorough consideration of potential risks, i.e., Risk-related Decision-Making (RDM) (Dalley and Robbins, 2017). While moderate impulsive action and RDM are considered normal personality traits, excessive impulsivity may contribute to psychiatric disorders such as drug abuse. Indeed, clinical research shows that psychostimulant abusers display heightened impulsive action (Voon et al., 2014) and RDM (Kjome et al., 2010) compared to healthy subjects, and in animals, those impulsivity facets predict cocaine intake (Dalley et al., 2007; Arrondeau et al., 2023) and relapse (Ferland and Winstanley, 2017), respectively. Given those findings, elucidating the neuronal circuits of impulsivity facets could help understand the individual vulnerability to drug abuse.

In an effort to identify the neural underpinnings of impulsivity, neuroimaging research in humans has evaluated the contribution of the medial prefrontal cortex (mPFC) function to specific impulsivity facets. Such research has shown that individuals with mPFC lesions display higher trait impulsivity (McDonald et al., 2017) and RDM (Clark et al., 2008) than healthy controls. Besides, in healthy populations, reduced mPFC activity has been associated with heightened trait impulsivity (Boes et al., 2009; Neal and Gable, 2017), impulsive action (Neufang et al., 2016) and RDM (Lv et al., 2021). However, despite the association between mPFC activity and impulsive behaviors in humans, preclinical studies examining the direct influence of the mPFC on impulsive action and RDM have yielded inconsistent results. For instance, in some studies, lesions or pharmacological inactivation of the infralimbic and prelimbic cortex (i.e., rodent homologs of the human mPFC) have been shown to increase impulsive action (Chudasama et al., 2003; Murphy et al., 2012; Feja and Koch, 2014) and bias choices towards disadvantageous options (Paine et al., 2013; Zeeb et al., 2015; Orsini et al., 2018; van Holstein and Floresco, 2020). However, in other studies, lesions (Paine et al., 2013), pharmacological (Zeeb et al., 2015), or chemogenetic (van der Veen et al., 2021) inhibition of the mPFC had no effect on impulsive action or RDM (Barrus et al., 2017). Given these discrepancies, further research is needed to clarify the contribution of the mPFC in modulating impulsive behaviors.

The mPFC might modulate impulsive action and RDM by exerting top-down control on subcortical structures (Dalley et al., 2011). Tracer studies (Carr and Sesack, 2000; Murugan et al., 2017; Souza et al., 2021) have shown that the mPFC sends direct projections to subcortical structures such as the dorsal striatum (DST), the nucleus accumbens (NAc), and the ventral tegmental area (VTA), which are critically involved in impulse control (Dalley and Robbins, 2017). For instance, studies in both humans and rodents have linked trait impulsivity (Buckholtz et al., 2010), impulsive action (Bellés et al., 2021; Urueña-Mendez et al., 2023), and RDM (Oswald et al., 2015) to heightened evoked dopamine (DA) release in the striatum. Additionally, in rodents, RDM has also been associated with elevated firing rates of VTA-DA neurons (Freels et al., 2020). Interestingly, the mPFC may modulate both striatal DA release (Hill et al., 2018) and VTA-DA neuronal activity (Tan et al., 2014; Jo and Mizumori, 2016). Thus, evidence suggests that the mPFC could modulate impulsivity through pathway-specific projections. However, few studies have explored the role of specific mPFC projections on impulsivity, focusing primarily on impulsive action. For instance, in one study (de Kloet et al., 2021), chemogenetic inhibition of the mPFC-to-DST, but not mPFC-to-NAc increased impulsive action, while contrasting results occurred in another study when inhibiting the mPFC-to-DST pathway (Nakayama et al., 2018). Critically, no studies have yet evaluated the role of the mPFC-to-VTA pathway in either impulsivity facet.

Here, we assessed the effect of chemogenetic manipulations of the mPFC-to-VTA pathway in controlling impulsive behaviors. We used the Roman High- and Low-avoidance rat lines, which respectively display high- and low-impulsive behaviors (Bellés et al., 2023), to first explore potential baseline differences in brain metabolic activity using in vivo positron emission tomography (PET) and the radiotracer [18F]-fluorodeoxyglucose ([^18^F]-FDG). Next, we employed an intersectional chemogenetic approach to assess the effects of targeted activation or inhibition of the mPFC-to-VTA pathway on impulsive action, RDM, and mPFC/VTA metabolic activity in RHA and RLA rats, respectively. Our results indicate a specific regulatory role for the mPFC-to-VTA pathway in impulsive action, but not in RDM, highlighting the need to further explore the contribution of this pathway in impulsivity-related disorders.

## Material and methods

### Subjects

Subjects were adult male rats RHA (n=18) and RLA (n=18) from our outbred colony at the University of Geneva. Rats were paired or trio housed in a temperature-controlled room (22 ±2°C) on a 12h light-dark cycle. Animals were maintained at 85% of their free-feeding weight during behavioral tasks and received water ad libitum. All the experiments were approved by the animal ethics committee of the Canton of Geneva and implemented according to the Swiss federal law on animal care.

### General procedure

**Figure 1. A** shows the timeline of the study. Rats were trained in the rat Gambling task (rGT) for approximately 24 days. Then, they underwent stereotaxic surgery to specifically target the mPFC-to-VTA pathway. Rats received a retrogradely expressing Cre recombinase virus in the VTA along with an anterograde Cre-dependent design receptor activated by a design drug (DREADD)-expressing virus in the mPFC. Specifically, high-impulsive/RHA rats (n=12) received an excitatory hM3Dq-DREADD, while low-impulsive/RLA rats (n=12) received an inhibitory hM4Di-DREADD. This approach allowed us to investigate whether line-specific targeting of the mPFC-to-VTA pathway modulates impulsive behaviors. Control animals expressing a Cre-dependent control protein instead of the DREADD were included for each rat line (n=6). After surgical recovery, rats were re-trained in the rGT and tested until stable performances were achieved. Rats then underwent testing sessions under saline vehicle and were subsequently tested under Clozapine-N-oxide (CNO), the DREADD-activating drug. Three days after the last rGT session, rats received two locomotion tests, one under saline and one under CNO. Those tests were separated for a week, and the drugs were administered in a counterbalanced order. Finally, rats received a positron emission tomography (PET) scan with [^18^F]-FDG to measure their brain metabolic activity under saline and CNO. Scans were conducted one week apart, and the drugs were counterbalanced between scans.

**Figure 1.**
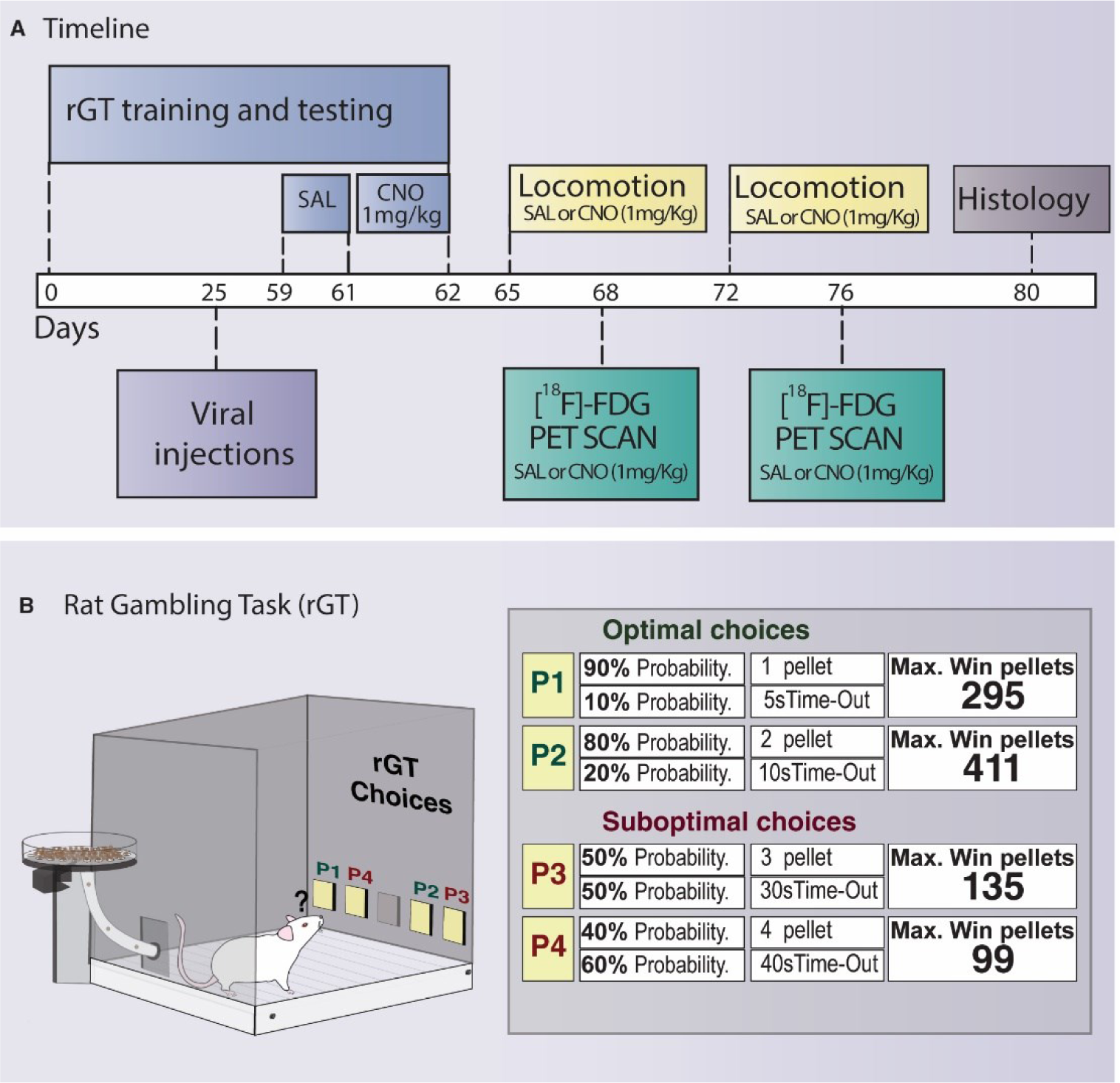
**A)** Timeline of the study with annotated time points (in days). **B)** rGT contingencies as described in Zeeb (2009).

### Drugs

CNO (HelloBio, Bristol, UK) was dissolved in 0.9% sterile saline at a concentration of 1mg/mL and administered intraperitoneally at 1mg/Kg 30min before testing. The vehicle solution consisted of 0.9% sterile saline injected at 1mL/Kg.

### Stereotaxic surgery and viral vectors

Details on surgical procedures are provided in the supplementary methods. Rats were anesthetized and placed in a stereotaxic frame (RWD Life Science, Mainz, Germany). A craniotomy was performed at the target coordinates [in the VTA: AP=0.37, ML=±0.09, and DV=-0.7mm; in the mPFC: AP=1.2, ML=±0.04, DV=-0.37mm]. The AP coordinate was taken from lambda, and the DV coordinate was taken from the dura. Viruses were delivered bilaterally (500nL/site) at a rate of 3nL/sec, using a nanoinjector (Nanoinject II, Drummond Scientific Company, Pennsylvania, USA). In the VTA, all animals received an AAV-hSyn-EGFP-Cre retrograde (8.8×1012 vg/mL, Viral Vector Facility, ETH, Switzerland). In the mPFC, viruses were infused as follows. For mPFC-to-VTA activation experiments, RHAs received AAV5-hSyn-DOI-hM3D(Gq)-mCherry (7×10¹²vg/mL, Addgene, plasmid number 44361). Conversely, for mPFC-to-VTA inhibition experiments, RLAs received AAV5-hSyn-DOI-hM4D(Gi)-mCherry (7×10¹²vg/mL, Addgene, plasmid number 44362). As a control, a subgroup of animals within each rat subline received AAV5-hSyn-DOI-mCherry (7.10¹² vg/mL, Addgene, plasmid number 50459). After injections, the skull was sutured and animals were allowed to recover for one week. Animals received pre- and post-operative analgesia and were fed ad-libitum until behavioral testing resumed.

### Rat Gambling Task (rGT)

A detailed rGT training and testing protocol is presented in the supplementary methods. In brief, rats were trained in daily 30min sessions as specified in Zeeb (2009). Animals learn to nose-poke into four holes termed P1, P2, P3, and P4 (**Fig.1B**). The holes varied in the probability of receiving a specific number of food pellets or a time-out (TO) punishment of varying duration. On rewarded trials, rats received the specified number of pellets. On punished trials, TO was signaled by a light flashing at 0.5Hz in the selected hole. P1 and P2 options were considered optimal, as they resulted in more gains and less punishment in each session. Conversely, the P3 and P4 were considered suboptimal as they resulted in fewer gains and more penalties in each completed session. Each trial was separated by an intertrial interval (ITI) of 5sec, responses during this interval were considered premature. rGT testing continued until the choice behavior stabilized (i.e., having ≤25% variation in the choice score during three consecutive days). Next, animals were i.p. injected with sterile saline 0.9% (1mL/Kg) 30min before testing for three days. The average of those sessions served as a comparison baseline. The following day, rats received an i.p. injection of CNO (1mg/Kg) 30min before testing.

A choice score [(%P1+%P2)–(%P3+%P4)] was calculated as an index of risk-related decision-making (Zeeb and Winstanley, 2013; Fugariu et al., 2020), with a lower choice score indicating less optimal decision-making. Impulsive action was measured as the percentage of premature responses performed during the ITI (i.e., number of premature responses/total number of trials initiated x 100; Ferland and Winstanley, 2017).

### [^18^F]-FDG PET scan

#### [^18^F]-FDG image acquisition and reconstruction

Rats underwent 2 PET scans, one under saline and one under CNO. Scans were performed one week apart and the treatments were administered in a counterbalanced order. Animals were fasted for 12 hours before each scan to minimize competition between dietary glucose and the [^18^F]-FDG radiotracer (Ribeiro et al., 2022). Rats then received an ip. injection of CNO (1mg/Kg/mL) or an equal volume of saline vehicle. Next, rats were implanted with a polyurethane catheter in the tail vein and placed in polystyrene boxes for 30min until receiving an intravenous bolus injection of 40 ± 5 MBq of [^18^F]-FDG (Department of Nuclear Medicine, Geneva University Hospitals). The doses injected were not significantly different between rat lines or Saline/CNO conditions (RHA: SAL=39.07±4.52 MBq and CNO=38.34±6.38 MBq; RLA SAL=41.08±4.65 MBq and CNO=40.50±4.11 MBq; Line: F_(1,33)_=1.70, p=0.20; Time: F_(1,33)_=1.36, p=0.25; Line × Time: F_(1,33)_=0.018, p=0.89). After [^18^F]-FDG injection, rats were placed in standard home cages and kept warm with a red-light lamp to prevent brown tissue uptake of the radiotracer (Ribeiro et al., 2022). At 45min uptake, anesthesia was initiated and maintained with isoflurane (2.5%) in oxygen, and the rats were positioned by two in the micro-PET scanner Triumph II (TriFoil Imaging, Northridge, CA, USA) using a compatible custom-made stereotaxic-like frame. At 50min post-radiotracer injection, animals were scanned for 20min. Dynamic PET images were acquired in list mode and reconstructed into 4-time frames of 5min each using the ordered subset expectation maximization algorithm with 20 iterations.

#### [^18^F]-FDG analysis

Specific [^18^F]-FDG brain templates and regions of interest (ROIs) were created for each rat line using PMOD software (version 4, PMOD Technologies Ltd., Zurich, Switzerland) as described in the supplementary methods. Individual PET scans were then converted into SUV units according to the formula: [R / (A/W)], where R is the radioactivity concentration (kBq/cc), A is the decay-corrected amount of injected radiotracer, and W is the weight of the rat in grams. Each individual PET scan was then averaged to obtain corresponding PET summation images. Individual PET summation images were coregistered to the specific RHA or RLA brain template using mutual information-based rigid body registration, and the resulting transformation was applied to the corresponding PET dynamic images. PET dynamic images were then normalized using a whole-brain normalization factor (NF=Average whole-brain uptake of all animals per group/individual whole-brain uptake), as described in Casado-Sainz et al., (2022). A specific template containing the following ROIs: medial prefrontal cortex (mPFC), midbrain, orbitofrontal cortex (OFC), cingulate cortex (Cg) dorsal striatum (DST), and ventral striatum (VST) was then applied to the resulting normalized PET images, and the normSUV values were extracted per each region.

### Locomotor activity

Locomotor testing occurred in four identical 48×48×40cm transparent open field boxes (ActiMot, TSE Systems, Bad Homburg, Germany). Tests were performed one week apart, once under saline vehicle and once under CNO (1mg/Kg). Treatments were administered in a counterbalanced order 30 min before testing and the total traveled distance was recorded for 30min.

### Tissue preparation and histology

Rats were anesthetized using sodium pentobarbital (150 mg/Kg, ip., at 200mg/mL) and transcardially perfused with 4% paraformaldehyde (PFA) in 0.1M phosphate-buffered saline (pH=7.4). Brains were extracted and stored in 4% PFA overnight at 4 °C and then transferred to a 30% sucrose solution at 4°C for 48-72h before being snap-frozen in isopentane at -55°C. Brains were sliced into 40-μm coronal sections using a cryostat (CM3050, Leica Biosystem, Muttenz, Switzerland) and stored in a cryoprotectant solution at −20°C for further processing. Free-floating sections were washed three times (10 min each) in PBS 0.1M and then stained with Hoechst-33342 dye (1:1000μl) for 30min at room temperature. Stained sections were washed three times per 10min each and mounted on slides covered with antifade mounting medium (Fluorsave, Merck, Darmstadt, Germany). Stained sections were imaged at 10X using a widefield fluorescence slide scanner microscope (Zeiss Axioscan Z1, Gottingen, Germany).

### Statistical analysis

All statistical analyses were conducted with SPSS Statistics 26.0 (IBM) software. The normality of data distribution was verified using the Shapiro-Wilk statistic at p<0.05, and for ANOVA analysis, non-normal data were LOG-10 transformed, as specified below.

Baseline behavioral differences between rat lines (i.e., RHA vs. RLA rats under saline) were analyzed with unpaired t-tests or Mann-Whitney-U statistics, depending on the data distribution. Between line differences in [^18^F]-FDG normSUV values at baseline were analyzed using a repeated measures ANOVA with rat line as between-subject factor and brain region as within-subject factor. Upon violation of data sphericity, the Greenhouse–Geisser correction was applied. The effect of mPFC-to-VTA pathway manipulation on behavioral measures was analyzed for each rat line using a repeated measures ANOVA, with Virus (i.e., DREADD or mCherry) as a between-subjects factor and treatment (i.e., saline or CNO) as a within-subjects factor. An additional analysis of the percentage of change under CNO was performed using unpaired t-tests or Mann-Whitney-U statistics when appropriate. The effects of mPFC-to-VTA pathway manipulation on [^18^F]-FDG normSUV values in the mPFC/midbrain regions were evaluated with a mixed factorial ANOVA with Virus (i.e., DREADD or mCherry) as a between-subjects factor, and treatment (i.e., saline or CNO) and brain region as within-subjects factors. Due to a technical issue, the saline PET images for one RLA rat were lost, so this animal was excluded from the PET analysis.

## Results

### Baseline differences in impulsive behaviors, locomotor activity and brain [^18^F]-FDG uptake in Roman rats

At baseline, RHA rats committed more premature responses (t_(36)_=6.4, df=34, p<0.0001, d=2.15; **Fig. 2A**) and reached lower choice scores than RLA rats (U_(36)_=94, p=0.031, η^2^=0.13; **Fig. 2B**), indicating that RHA rats display greater impulsive action and less optimal decision-making, respectively. Although both rat lines initiated a comparable number of trials (t_(36)_=-0.83, df=34, p=0.41, d=0.04; **Fig. 2C**), RHA rats committed fewer omissions (t_(36)_=-2.66, df=34, p=0.01, d=0.90; **Fig. 2D**), suggesting that RHA rats have a higher motivation for the task. We also observed higher baseline locomotor activity in RHA than in RLA rats; however, this difference did not reach statistical significance (U_(36)_=107, p=0.09, η^2^=0.08; **Fig. 2E**).

**Figure 2.**
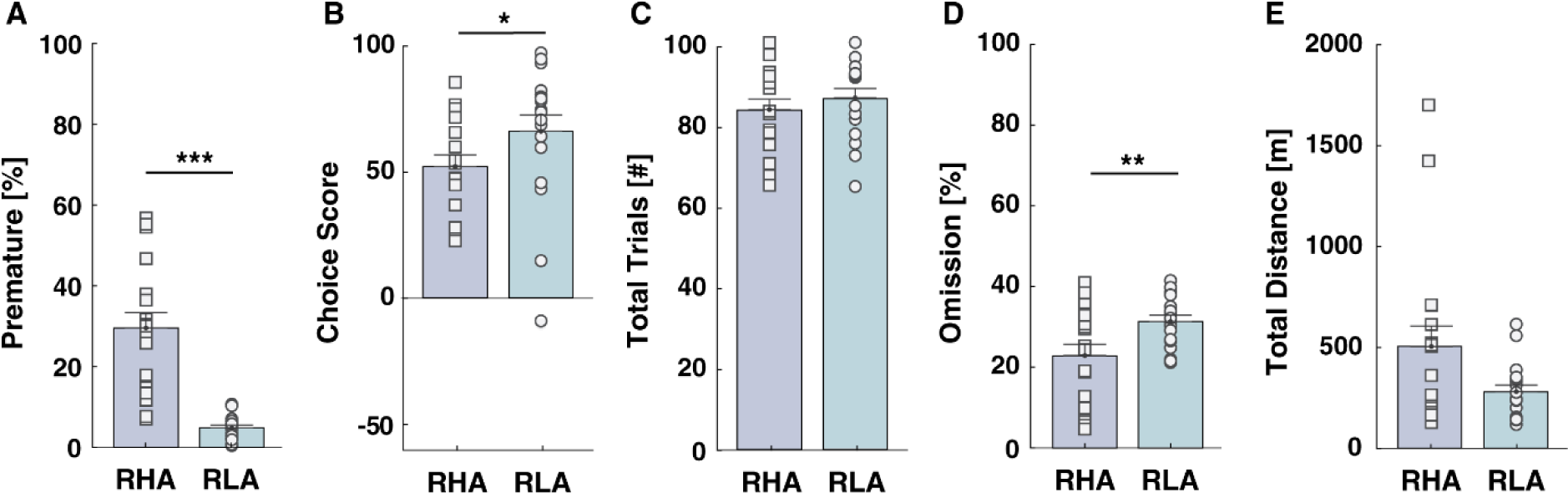
Baseline rGT performance and locomotor activity in Roman rats. RHA rats **A)** display a higher percentage of premature responses and **B)** reach lower choice scores than RLA rats. **C)** Both rat lines initiated a comparable number of trials. However, **D)** RHA rats displayed a lower percentage of omissions. **E)** No significant difference occurred in the total traveled distance during the locomotion test. However, RHA rats tended to travel greater distances than RLAs.

Mean parametric maps for baseline [^18^F]-FDG normSUV values in RHA and RLA rats are presented in **Figure 3A**. As observed, RHA rats displayed lower [^18^F]-FDG normSUV values compared with RLA rats. Such differences were observed along cortico-midbrain-striatal structures. There was a main effect of Line (F_(1,33)_= 268, p<0.001, ηp^2^=0.99) and a Line x Brain region interaction (F_(3.5,115)_= 2.94, p<0.05, ηp^2^=0.08) on [^18^F]-FDG normSUV values, which was significant for all evaluated brain regions (p<0.001). These results indicate that RHA rats display lower metabolic brain activity than RLA rats.

**Figure 3.**
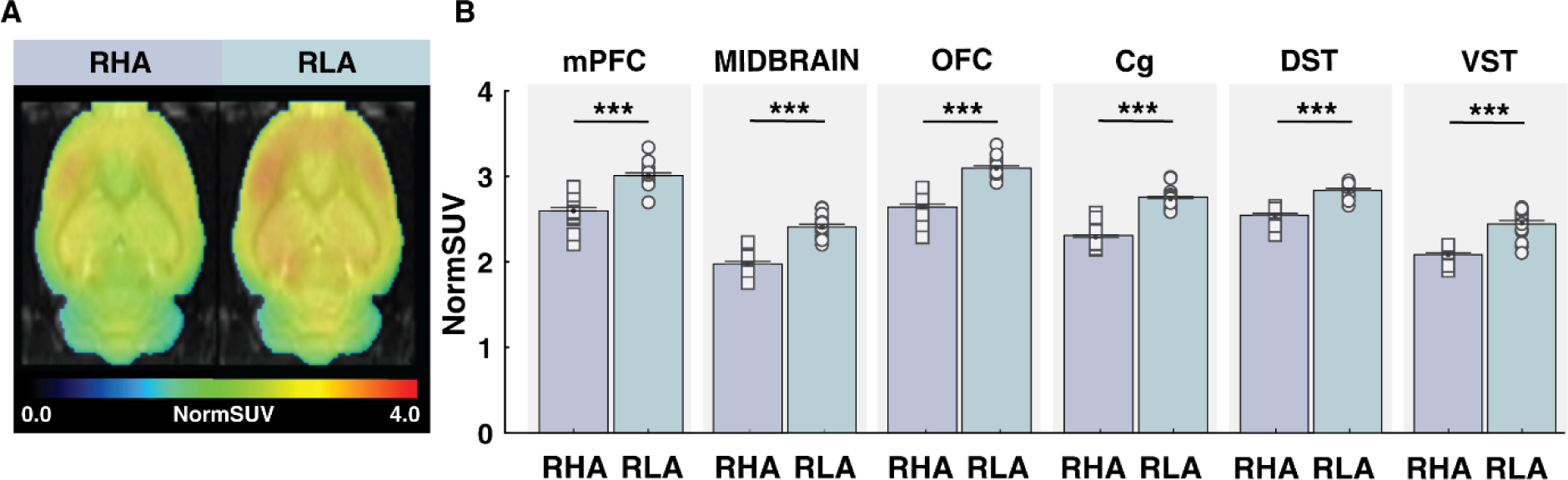
Baseline [^18^F]-FDG uptake in Roman rats. **A)** Mean parametric maps of [^18^F]-FDG normSUV in RHA and RLA rats. Normalized normSUV maps are projected upon the MRI rat atlas (greyscale) and are shown in horizontal planes at the level of the mPFC. **B)** RHA rats displayed lower [^18^F]-FDG uptake than RLA rats in the mPFC, Midbrain, OFC, Cg, DST, VST and VST**. [^18^F]-FDG:** [18]Fluorodeoxyglucose. **mPFC**: medial Prefrontal Cortex, **OFC**: orbitofrontal Cortex, **Cg**: Cingulate Cortex, **DST:** Dorsal Striatum, **VST:** Ventral Striatum.

### The mPFC-to-VTA pathway modulates impulsive action but not RDM

A representative image of the viral expressions in the mPFC is presented in **Figure 4A**. Manipulations of the mPFC-to-VTA pathway significantly affected the percentage of premature responses (**Fig 4. B-E**). In RHA rats, we observed a significant Virus x Treatment interaction (F_(1,16)_=7.25, p<0.05, ηp^2^=0.31), but no main effects of virus (F_(1,16)_=0.22, p=0.64, ηp^2^=0.01) or treatment (F_(1,16)_=4.28, p=0.06, ηp^2^=0.2). Post-hoc comparisons revealed that CNO reduced premature responding in hM3Dq-expressing RHAs (p<0.001) but not in controls (p>0.05). An additional analysis of the percentage of change under CNO relative to saline further confirmed this result (hM3Dq-expressing vs control RHA rats: t_(18)_=4.01, df=16, p<0.01, d=1.92), suggesting that the reduction in premature responding was specific to the mPFC-to-VTA pathway activation and not to nonspecific effects of CNO.

**Figure 4.**
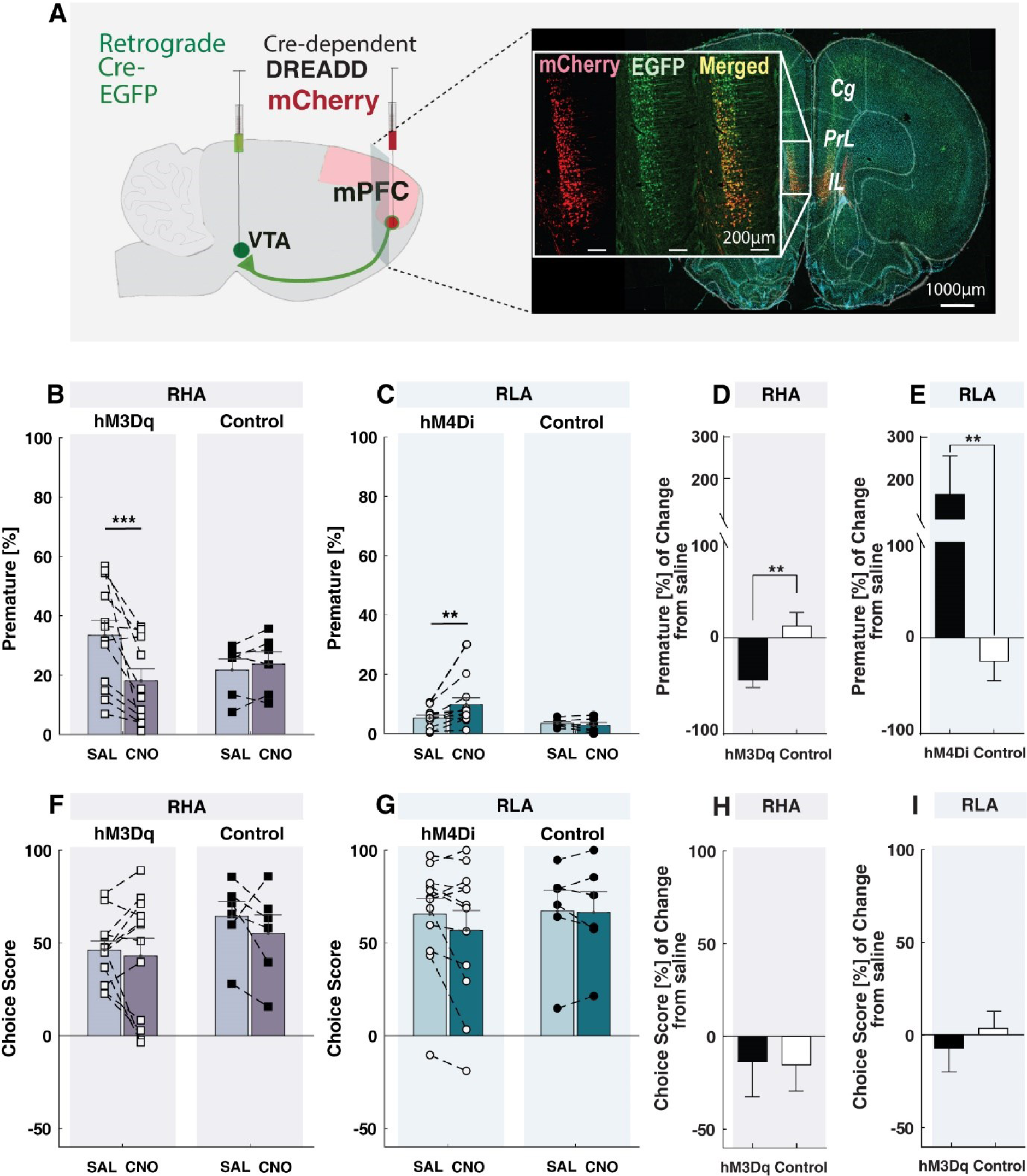
Chemogenetic manipulation of the mPFC-to-VTA pathway on impulsive action and RDM. **A)** Fluorescent microscopy of the virally transduced mCherry in the mPFC. **B)** Chemogenetic activation of the mPFC-to-VTA pathway in RHA rats decreased premature responding. **C)** Conversely, chemogenetic inhibition of the mPFC-to-VTA pathway in RLA rats increased premature responding. Similar results were observed when measuring the percentage of change in premature responses after CNO in **D)** RHA and **E)** RLA rats. Chemogenetic manipulation of the mPFC-to-VTA pathway did not alter the choice score in either **F)** RHA or **G)** RLA rats. Similar negative results were observed when measuring the percentage of change in choice score after CNO in **H)** RHA and **I)** RLA rats.

In RLA rats, there was also a significant Virus x Treatment interaction (F_(1,16)_=4.68, p<0.05, ηp^2^=0.23), but not main effects of virus (F_(1,16)_=3.72, p=0.07, ηp^2^=0.19) or treatment (F_(1,16)_=2.53, p=0.13, ηp^2^=0.14). Post-hoc comparisons revealed that CNO increased premature responding in hM4Di-expressing RLAs (p<0.001) but not in controls (p>0.05). This result was further confirmed by an additional analysis of the percentage of change in premature responding under CNO relative to saline (hM4Di-expressing vs control RLA rats: U_(18)_=6, p<0.01, η^2^=0.46), indicating that inhibition of the mPFC-to-VTA pathway, rather than unspecific effects of CNO, increased premature responding. Altogether, our results suggest that activation of the mPFC-to-VTA pathway specifically reduces impulsive action, while inhibition has the opposite effect.

When evaluating the effect of mPFC-to-VTA pathway manipulations on RDM (**Fig 4. F-I**), we observed no virus x treatment interaction in either rat line (RHA: F_(1,16)_=0.28, p=0.60, ηp^2^=0.02; RLA: F_(1,16)_=1.29, p=0.27, ηp^2^=0.07). There were also no main effects of virus (RHA: F_(1,16)_=1.83, p=0.19, ηp^2^=0.10; RLA: F_(1,16)_=0.13, p=0.72, ηp^2^=0.01), nor main effect of treatment in any rat line (RHA: F_(1,16)_=1.13, p=0.30, ηp^2^=0.07; RLA: F_(1,16)_=1.84, p=0.19, ηp^2^=0.10), suggesting that the activity of the mPFC-to-VTA pathway does not control RDM. Analysis of the percentage change in RDM under CNO, relative to saline treatment, further revealed no significant differences between the DREADD-expressing and control groups in either rat line (RHA: t_(18)_=0.07, df=16, p=0.94, d= 0.03; RLA: t_(18)_=0.64, df=16, p=0.5, d=0.36). Altogether, our results suggest that the mPFC-to-VTA pathway specifically modulates impulsive action but not RDM.

### Chemogenetic manipulation of the mPFC-to-VTA pathway on other behavioral measures

Neither activation nor inhibition of the mPFC-to-VTA pathway affected the total number of trials initiated in the rGT (**Fig. 5A-D**) in either rat line (virus x treatment in RHA: F_(1,16)_=0.31, p=0.58, ηp^2^=0.02; in RLA: F_(1,16)_=0.04, p=0.84, ηp^2^=0). There was also no main effect of virus (in RHA: F_(1,16)_=0.91, p=0.36, ηp^2^=0.02; in RLA: F_(1,16)_=0.50, p=0.49, ηp^2^=0.03) nor effect of treatment (in RHA: F_(1,16)_=0.31, p=0.58, ηp^2^=0.02; in RLA: F_(1,16)_=0.37, p=0.55, ηp^2^=0.02).

**Figure 5.**
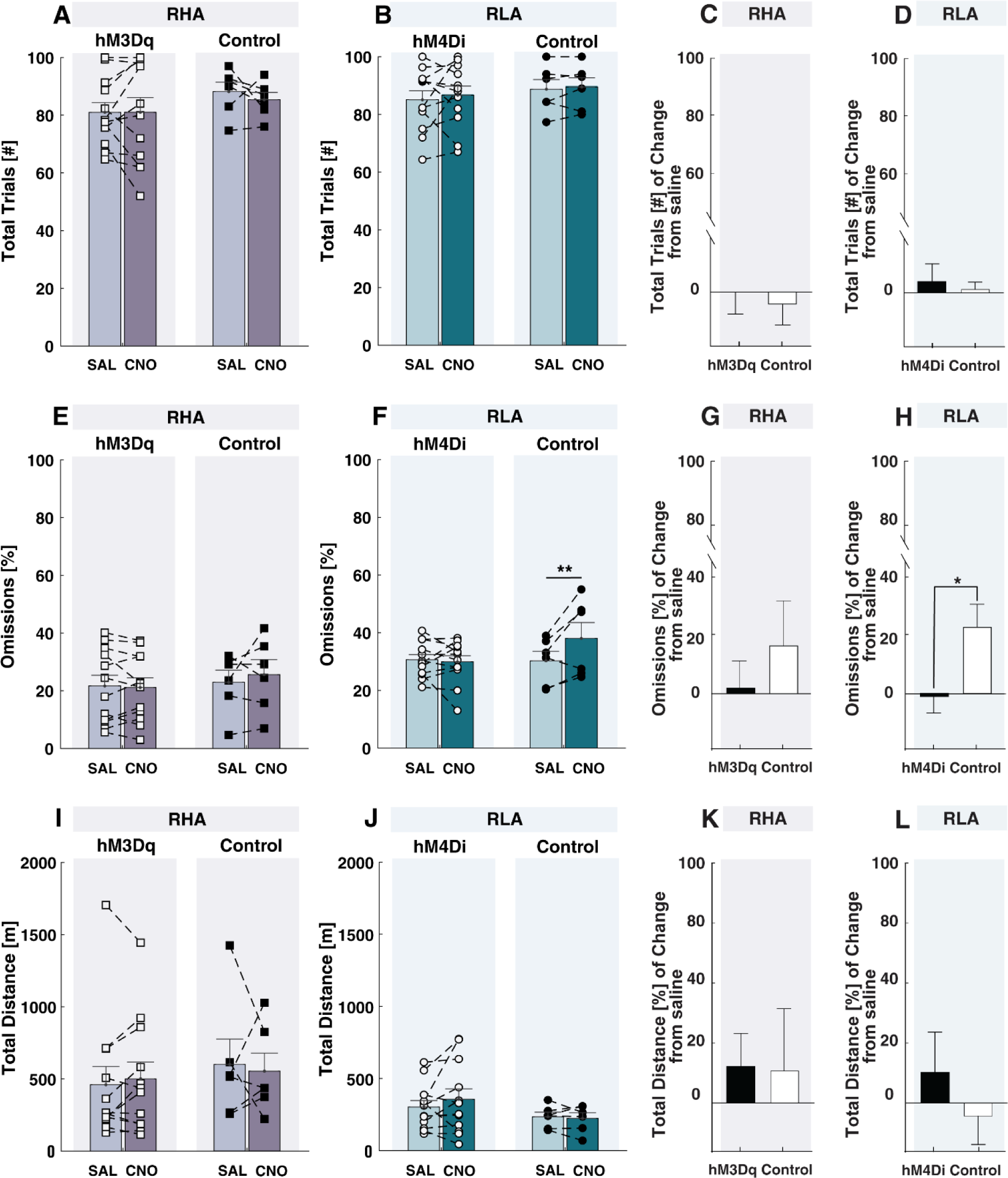
Chemogenetic manipulation of the mPFC-to-VTA pathway on other behavioral measures. Chemogenetic manipulation of the mPFC-to-VTA pathway in RHA or RLA rats did not significantly alter **A-D)** the total number of trials initiated and **E-H)** the percentage of omissions in the rGT, or **I-L)** the locomotor activity in an open field as measured by the total distance traveled.

These findings were consistent with an analysis of the percentage of change under CNO, relative to saline treatment, where no significant differences occurred between DREADD-expressing and control animals in either rat line (In RHA rats: t_(18)_=0.35, df=16, p=0.72, d=0.18; in RLA rats: t_(18)_=0.31, df=16, p=0.76, d=0.18).

**Figure 5. E-H** shows the effect of the mPFC-to-VTA pathway manipulations on the percentage of omissions. In RHA rats, we observed no Virus x Treatment interaction (F_(1,16)_=1.05, p=0.32, ηp^2^=0.06), nor main independent effect of virus (F_(1,16)_=0.25, p=0.63, ηp^2^=0.02) or treatment (F_(1,16)_=0.46, p=0.51, ηp^2^=0.03). There was also no significant percentage of change under CNO relative to saline between hM3Dq-expressing and control RHA rats (t_(18)_=0.84, df=16, p=0.41, d=0.40). Conversely, in RLA rats, we observed a virus x treatment interaction (F_(1,16)_=7.43, p=0.01, ηp^2^=0.32) and treatment effect (F_(1,16)_=5.10, p=0.04, ηp^2^=0.4) on omissions. Post-hoc analyses revealed that omissions were higher only in RLAs control after CNO (p<0.01). The percentage of change in omissions in CNO relative to saline treatment was significantly increased only in RLA controls (t_(18)_=2.61, df=16, p<0.05, d=0.13). This result might be secondary to a higher variability and lower n within this control group.

Finally, neither activation nor inhibition of the mPFC-to-VTA pathway affected locomotion activity (**Fig. 5. I-L**). We observed no virus x treatment interaction (RHA: F_(1,16)_=2.44, p=0.63, ηp^2^=0.02; RLA: F_(1,16)_=0.22, p=0.69, ηp^2^=0.01), or main effects of virus (RHA: F_(1,16)_=0.53, p=0.37, ηp^2^=0.05; RLA: F_(1,16)_=0.66, p=0.43, ηp^2^=0.04) or treatment (RHA: F_(1,16)_=0.05, p=0.82, ηp^2^=0.03; RLA: F_(1,16)_= 0.16, p=0.69, ηp^2^=0.01) in either rat line. Analysis of the percentage of change in locomotion under CNO relative to saline treatment showed no significant differences between DREADD-expressing and control animals in either rat line (In RHA: t_(18)_=0.08, p=0.94, d=0.04; in RLA: t_(18)_=0.86, df=16, p=0.4, d=0.1). Taken together, our results further suggest that the effects of the mPFC-to-VTA manipulations on impulsive action are not related to changes in locomotor functions.

### Chemogenetic manipulation of the mPFC-to-VTA pathway does not lead to detectable changes in [^18^F]-FDG uptake

**Figure 6** presents [^18^F]-FDG NormSUV values for the mPFC and midbrain after chemogenetic manipulations. Chemogenetic activation or inhibition of the mPFC-to-VTA pathway led to no virus x treatment x structure interaction in RHA or RLA rats, respectively (RHA: F_(1,16)_=0.5 p=0.49, ηp^2^=0.03; RLA: F_(1,15)_=0.33, p=0.57, ηp^2^=0.02), nor main effects of virus (RHA: F_(1,16)_=0.15, p=0.71, ηp^2^=0.01; RLA: F_(1,15)_=0.13, p=0.73, ηp^2^=0.01) or treatment (RHA: F_(1,16)_=0.03, p=0.87, ηp^2^=0; RLA: F_(1,15)_=1.32, p=0.27, ηp^2^=0.08). Altogether, our results suggest that chemogenetic manipulations of the mPFC-to-VTA pathway resulted in no detectable changes in [^18^F]-FDG uptake within the mPFC or the midbrain.

**Figure 6.**
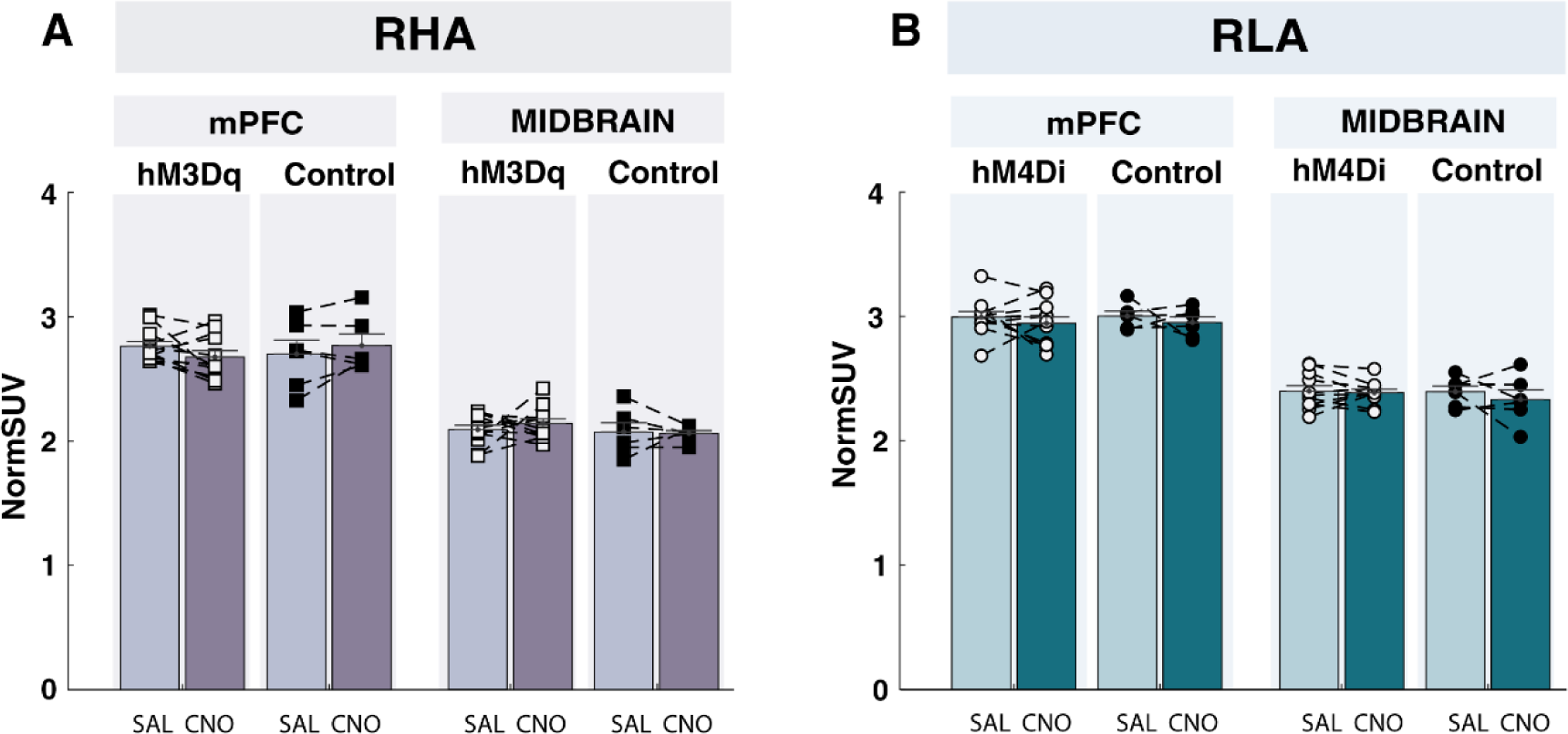
Chemogenetic manipulation of the mPFC-to-VTA pathway on [18F]-FDG uptake. No alterations in [^18^F]-FDG uptake were detected in the mPFC or midbrain after chemogenetic manipulation of the mPFC-to-VTA pathway in either **A)** RHA or **B)** RLA rats.

## Discussion

Our study offers new insights into the underlying mechanisms of impulse control by indicating a direct contribution of the mPFC-to-VTA pathway in controlling a specific impulsivity facet. Extending previous studies (Arrondeau et al., 2023; Bellés et al., 2023) showing greater impulsive action and less optimal decision-making in RHA than RLA rats, we evidenced reduced cortical [^18^F]-FDG uptake in the more impulsive phenotype. This result strengthens the view that reduced cortical function could be an underlying alteration to impulsive behaviors. Importantly, by specifically targeting the mPFC-to-VTA pathway in a phenotype-dependent way, we were able to revert innate patterns of impulsivity. Chemogenetic activation of the mPFC-to-VTA pathway in high-impulsive RHA rats reduced premature responding, while chemogenetic inhibition of this pathway in low-impulsive RLA rats had the opposite effect. Our findings indicated that the mPFC-to-VTA pathway selectively controls impulsive action, as neither chemogenetic manipulation altered RDM, further suggesting that both impulsivity facets, while related, depend on separate neural circuits. Nevertheless, despite a robust behavioral effect on impulsive action, changes in brain [^18^F]-FDG uptake were undetectable. Taken together, our results support a regulatory role for the mPFC-to-VTA pathway on impulsive action, suggesting a potential target for understanding impulsivity-related disorders.

Our observation of lower brain [^18^F]-FDG uptake in RHA than RLA rats at baseline complements prior studies suggesting reduced glucose metabolism in the cingulate (Cg) and orbitofrontal cortex (OFC) of highly impulsive animals (Barbelivien et al., 2001). Moreover, our results extend previous studies showing that RHA rats exhibit reduced frontocortical volume (Río-Álamos et al., 2019) and decreased cortical glutamate receptors than RLA rats (Klein et al., 2014), supporting the idea that reduced frontocortical function might be linked to heightened impulsivity. Together with these reports, our observation of cortical hypometabolism in impulsive rats parallelled human studies, where individuals with high self-reported impulsivity (Matsuo et al., 2009; Neal and Gable, 2017) and RDM (Aydogan et al., 2021) exhibit reduced frontocortical activity and reduced cortical volume. Altogether, our results strengthen the view of frontocortical hypofunction as an alteration underlying heightened impulsivity.

Although the medial prefrontal cortex (mPFC) exerts a top-down control on impulsivity-related structures (Dalley et al., 2011), including the VTA, no previous study explored the role of the mPFC-to-VTA pathway in the control of impulsive behaviors. By using a within-subjects design and a chemogenetics strategy suited to the rats’ impulsive phenotype, our study shows for the first time that the mPFC-to-VTA pathway modulates a specific impulsivity facet. Activating the mPFC-to-VTA pathway in RHA rats reduced their impulsive action, while inhibiting this pathway in RLA rats has the opposite effect. Notably, the absence of changes in overall locomotor activity after chemogenetic manipulations highlights the specificity of the mPFC-to-VTA pathway in the control of impulsive action. Our results are consistent with recent chemogenetic and optogenetic studies where unspecific activation of the mPFC improved impulsive action (Warthen et al., 2016; Suri et al., 2023), while unspecific inhibition impaired it (Luchicchi et al., 2016; Nakayama et al., 2018). Interestingly, our results extend those findings by indicating that the mPFC exerts a significant influence on impulsive action through its direct projections onto the VTA.

Impulsive action has been linked to dopaminergic function in the VTA (Hynes et al., 2021). High self-reported impulsivity in humans (Buckholtz et al., 2010) and heightened impulsive action in rodents (Bellés et al., 2021) have both been associated with reduced availabilities of dopamine (DA) type 2 autoreceptors in the VTA and increased evoked-DA release in the striatum. Moreover, optogenetic stimulation of the VTA-DA projections to the nucleus accumbens (NAc) increases impulsive action (Flores-Dourojeanni et al., 2021), while chemogenetic inhibition of these projections -at least in males-reduces impulsivity (Hynes et al., 2021). Thus, it is possible that our manipulations of the mPFC-to-VTA pathway controlled impulsive action by affecting VTA-DA activity. Although the exact mechanism of mPFC modulation on VTA-DA activity remains unclear, studies have shown that mPFC activity can exert both excitatory and inhibitory effects on VTA-DA neurons (Tong et al., 1996). For instance, electrical stimulation of the mPFC increases VTA-DA firing (Lodge, 2011) and DA release in the NAc (Taber et al., 1995), but it can also inhibit a subset of VTA-DA neurons (Tong et al., 1996; Gao et al., 2007). This observation is consistent with pharmacological studies where the mPFC inhibition increased VTA-DA firing (Patton et al., 2013; Jo and Mizumori, 2016). Interestingly, some studies proposed that mPFC activity inversely controls VTA-DA activity through direct activation of VTA-GABA neurons (Tan et al., 2014). Considering that the mPFC directly innervates VTA-GABA neurons (Carr and Sesack, 2000; Sesack and Grace, 2010), such GABAergic control could be a potential circuit for the mPFC-to-VTA pathway regulation of impulsive action. This hypothesis aligns with a recent study suggesting that projections from cortical structures (i.e., the cingulate) to the VTA could inhibit DA neurons via activation of VTA-GABA interneurons (Song et al., 2024). However, future work is needed to confirm whether this cellular mechanism underpins the mPFC-to-VTA pathway effects on impulsive action.

Our results indicate that while the mPFC-to-VTA pathway modulates impulsive action, it does not appear to influence RDM. This selective control of the mPFC activity across different facets of impulsivity is consistent with previous studies using a nonspecific mPFC inactivation method (Paine et al., 2013; Zeeb et al., 2015), suggesting that impulsive action and RDM may rely on overlapping yet distinct neuronal circuits. One possibility is that the mPFC-to-VTA pathway requires the convergent actions of other subcortical structures to control RDM. The rostromedial tegmental nucleus (RMTg), known to inhibit VTA-DA neurons and control risk-reward seeking (Vento et al., 2017), is one potential candidate. Differential activity of the RMTg between high and low-impulsive rats could counteract any potential effect of the mPFC-to-VTA pathway manipulation on RDM. Alternatively, the mPFC might recruit different projections to influence RDM such as those innervating the NAc or the basolateral amygdala (Orsini et al., 2015; Truckenbrod et al., 2023). Further investigation targeting additional subcortical structures or mPFC projections would help determine the precise underlying networks controlling RDM.

Chemogenetic manipulation of the mPFC-to-VTA pathway modulates impulsive action without inducing detectable changes in glucose metabolism. This result contrasts with previous research demonstrating detectable changes in brain glucose metabolism following chemogenetic manipulation of the nigrostriatal pathway (Casado-Sainz et al., 2022). One potential explanation for these differences is the relative density of projections within each specific pathway. While a precise quantification is currently lacking, it is reasonable to estimate that less than 20% of the mPFC neurons project to the VTA (Choi et al., 2024), compared to over 70% of the substantia nigra DA neurons projecting to the striatum (Chen et al., 2022). Thus, greater projection densities within the nigrostriatal pathway might have contributed to stronger/detectable brain metabolic changes. Altogether, our results suggest that while chemogenetic manipulation of the mPFC-to-VTA pathway influences impulsive action, its effect on brain glucose metabolism may have lied below the detection threshold of the PET scanning.

Our study opens the possibility that alterations within the mPFC-to-VTA pathway in specific phenotypes might play a role in impulsivity-related disorders such as drug abuse. Individuals with drug abuse disorders exhibit heightened impulsive action (Voon et al., 2014), frontocortical hypofunction (Volkow et al., 1992; Chen et al., 2007; Kim et al., 2009; Harlé et al., 2014), and disruptions in DA release (Martinez et al., 2007) compared to healthy controls. Interestingly, recent studies in rodents demonstrated that rats vulnerable to alcohol abuse (Gozzi et al., 2013) or compulsive cocaine seeking (Gozzi et al., 2011; Cannella et al., 2017; Jones et al., 2023) exhibit baseline alterations within cortical, striatal, and midbrain regions, consistent with those reported in impulsive-RHA rats. In line with these studies, our prior work indicates that high-impulsive RHA rats are more vulnerable to the rewarding effects of cocaine (Dimiziani et al., 2019) and to cocaine-related alterations of dopamine function than low-impulsive RLA rats (Urueña-Mendez et al., 2023). Thus, one interesting possibility is that the mPFC-to-VTA pathway activity not only controls impulsivity but lies at the intersection between impulsivity, frontocortical hypofunction, and individual vulnerability to drug abuse.

### Study limitations and future research

We used [^18^F]-FDG PET as a non-invasive indirect measure of large-scale brain activity. However, future studies might consider using additional molecular techniques, such as in vivo calcium imaging, which could provide a more sensitive measure of cortical activity in high- and low-impulsive rats during concurrent behavioral testing. Additionally, while we hypothesized that the mPFC-to-VTA pathway might exert its effects by acting on VTA-GABA neurons, confirming the VTA neuronal identity involved in this pathway would be necessary to support this hypothesis further. Finally, a growing body of research suggests that impulsivity-underlying mechanisms might be different in females than males (Hynes et al., 2021; Orsini et al., 2022). Thus, studies extending our work by including female subjects are essential to determine whether the mPFC-to-VTA pathway similarly controls impulsive action across sexes.

## Conclusion

Our study addresses a critical gap in evaluating the role of the mPFC-to-VTA pathway in regulating specific impulsivity facets. Using two rat lines with innate differences in impulsivity, we observed a hypocortical metabolism in the high-impulsive phenotype. Importantly, phenotype-specific chemogenetic manipulations of the mPFC-to-VTA pathway modulated impulsive behaviors. Our findings propose a direct role of the mPFC-to-VTA pathway in controlling impulsive action but not RDM, indicating that these impulsivity facets are governed by distinct mechanisms. While more sensitive methods are required to determine the impact of the mPFC-to-VTA pathway manipulations on cortical activity, our study suggests a promising new target for investigating impulsivity-related disorders.

## Supplementary methods

### Stereotaxic surgery and viral vectors

Rats were anesthetized with isoflurane in oxygen (3% during induction and 1% during maintenance) and placed on a heating pad in a stereotaxic frame (RWD Life Science, San Diego, USA). Their scalp skin was retracted to expose the skull and craniotomy was made at target coordinates in the VTA, [AP: 0.37, ML:±0.09 and DV:-0.7mm] and the mPFC [AP:1.2 ML: ±0.04 DV:-0.37mm]. In both cases, the AP coordinate was taken from lambda, and the DV coordinate was taken from the dura. Viruses were delivered bilaterally (500nL per site) at a rate of 3nL per second, using a pulled glass capillary controlled by a nanoinjector (Nanoinject II, Drummond Scientific Company, Pennsylvania, USA). Before injecting, the capillary remained in the injection site for one minute. In the VTA, all animals received an AAV-hSyn-EGFP-Cre retrograde (8.8×1012 vg/mL, Viral Vector Facility, ETH, Switzerland). In the mPFC, viruses were infused as follows. For mPFC-to-VTA activation experiments, RHAs received AAV5-hSyn-DOI-hM3D(Gq)-mCherry (7×10¹²vg/mL, Addgene, plasmid number 44361). Conversely, for mPFC-to-VTA inhibition experiments, RLAs received AAV5-hSyn-DOI-hM4D(Gi)-mCherry (7×10¹²vg/mL, Addgene, plasmid number 44362). As a control, a subgroup of animals within each rat subline received AAV5-hSyn-DOI-mCherry (7.10¹² vg/mL, Addgene, plasmid number 50459). After injections, the capillary remained in place for 10 min before being slowly removed (∼ 1min). The skull was sutured and animals were allowed to recover for one week. Animals received pre and post-operatory analgesia and were fed adlibitum until resumed behavioral testing.

### Rat Gambling Task

The task was performed in eleven standard operant conditioning chambers (Med Associates Inc., St. Albans, VT, USA). Each chamber had a house light, a food tray, and five response holes (the central inactivated during testing). Each aperture was fitted with a cue light and infrared beams to detect nose-poke responses.

Rats were trained in daily 30 min sessions as specified in Zeeb (2009). Animals learned to nose-poke in the food tray to initiate each trial. After 5s of inter-trial interval (ITI), one hole was illuminated, and the rat had 5s to nose-poke into that hole to obtain a food pellet reward (purified rodent tablets of 45 mg, Test Diet, Sandown Scientific, UK). Upon learning the nose-poke response, the rGT options were introduced over eight forced-choice (FC) sessions (four before stereotaxic surgeries and four after-surgery recovery). The rGT options were termed P1, P2, P3, and P4. They varied in the number of pellets delivered (1, 2, 3, or 4, respectively), the probability of pellet delivery (0.9, 0.8, 0.5, or 0.4, respectively), the time-out (TO) punishment probability (0.1, 0.2, 0.5, or 0.6, respectively) and the TO punishment duration (5s, 10s, 30s, or 40s, respectively). Each option was associated with specific holes counterbalanced between animals. On rewarded trials, rats received the specified number of pellets. There was no reward on punished trials, and TO punished was signaled with a light flashing at 0.5Hz in the selected hole. During each FC session, each option was individually presented in a pseudorandom order. Following FC sessions, rats were tested in the rGT task, i.e., with the four options simultaneously available. Testing continued until the choice behavior stabilized (i.e., having ≤25% variation in the choice score during three consecutive days). Next, animals were i.p. injected with sterile saline 0.9% (1mL/Kg) 30 min before testing for three days. The average of those sessions served as a comparison baseline. The following day, rats received an i.p. injection of CNO (1mg/Kg) 30 min before testing.

#### [^18^F]-FDG template generation

[^18^F]-FDG brain templates for RHA and RLA rats were developed in PMOD software (version 4, PMOD Technologies Ltd., Zurich, Switzerland) according to a protocol previously described (Vállez Garcia et al., 2015). Individualize brain scans were obtained by cropping the double-rat dynamic PET scans. Next, individual PET dynamic images were converted into SUV units according to the formula [r / (aʹ/w)], where r is the radioactivity concentration (kBq/cc), aʹ is the decay-corrected amount of injected radiotracer, and w is the weight of the rat in grams. Per each rat line, 10 individual scans were selected and one representative scan was used as a reference. Individual scans were normalized into the space of the reference and then were averaged into a single PET scan. This averaged scan was duplicated and flipped from left to right, and was then subsequently averaged with its flipped duplicated to create a symmetrical voxel-wise averaged template.

#### Volumes of interest

3D Volumetric atlas for each RHA and RLA [^18^F]-FDG template was constructed based on the Schiffer’s magnetic resonance image (MRI) and volume of interest (VOI) atlas of the rat brain (Schiffer et al., 2006) as follows. First, the MRI atlas was coregistered to each rat template using mutual information-based rigid body registration. Next, the information outside the brain was masked in both the templates and the MRI. The masked MRI was then subjected to automatic elastic coregistration to each template dimension to obtain an adjusted MRI for each RHA and RLA template. The resulting transformations were independently applied to the VOI atlas so that the spatially transformed VOIs were specific for each rat line brain template.

A region of interest (ROI) was then defined on each adjusted MRIs using the VOI atlas for RHA and RLA rats as a reference. The ROI template included the following brain regions: the medial prefrontal cortex (mPFC, 1mm x 1.2mm oval), the midbrain (1.7 mm x 0.6mm oval), the orbitofrontal cortex (OFC, 1.5mm x 0.7 mm oval ), the cingular cortex (Cg, 1.3mm x 1mm oval), the dorsal striatum (DST, 1.8mm circle) and the ventral striatum (VST, 1.2 mm circle). ROIs were placed on the central planes of each structure to minimize the partial volume effect.

